# *In-silico* Exploration of Drug Leads Targeting Mycobacterial Ag85C

**DOI:** 10.64898/2026.09.26.754622

**Authors:** Katewad Raghavendra Singh, Anuradha, Chitta Suresh Kumar

## Abstract

This study aims to identify potential inhibitors of the Ag85C enzyme from Mycobacterium tuberculosis through primary docking, pharmacophore modelling, and ADME/pharmacokinetics analysis. Delamanid and Pretomanid emerged as the top hits in primary docking, demonstrating strong binding affinities and interactions with key active site residues. A robust pharmacophore model was developed, identifying essential features for binding and used to screen 4254 antituberculosis compounds, yielding three promising candidates: F0314-0040, F0398-0505, and F0349-2716. These compounds exhibited high binding affinities in major docking studies. ADME and pharmacokinetics analysis revealed favourable Drug likeness and absorption profiles, supporting their potential as effective antituberculosis agents. The integration of computational methods provided a comprehensive approach for rational drug design.

## Background

Tuberculosis (TB) continues to be the leading cause of mortality globally, due to the rise of *Mycobacterium tuberculosis* (*Mtb*) strains that are resistant to multiple drugs (MDR) **[1].** The emergence of these treatment-resistant strains has greatly complicated worldwide attempts to control tuberculosis, highlight the need for immediate progress in the field of drug research and development **[12].** As per the World Health Organization (WHO), regions with the largest burden of TB, such as India, China, and Russia, also have a particularly high prevalence of MDR-TB (Organization 2021). The global challenge presented by MDR-TB highlights the urgent need for innovative therapeutic approaches capable of surmounting medication resistance and treatment of the disease **[9;7].**

A significant challenge in the treatment of TB is the bacterium’s capacity to develop resistance to medications, frequently by mutations that modify drug targets or strengthen bacterial survival **[3].** This resistance has prompted to identify new therapeutic targets within the bacterium’s biological functions. Out of these options, the Antigen 85 (Ag85) complex, consisting of three proteins (Ag85A, Ag85B, and Ag85C), has become a very promising focus of attention. These proteins play a vital role in the production of the Mtb cell wall, which is essential for the bacterium’s ability to survive and cause disease **[23; 19].** The Ag85 complex is crucial for both cell wall formation and the bacterium’s interaction with the host immune system, making it a dual target for drug development and vaccination research **[6; 35].**

Computational drug design has revolutionized the process of identifying and optimizing new drugs against Mycobacterium tuberculosis. This allows the researchers to simulate the interactions between potential drugs and their targets at the molecular level **[18].** Techniques such as molecular docking, virtual high-throughput screening, and molecular dynamics simulations have become invaluable tools These methods enable the identification of compounds that could not only bind strongly to the target protein but also exhibit the desirable pharmacokinetic properties, such as stability and bioavailability. For instance, a study utilizing these techniques identified novel inhibitors that bind with high stability to the Ag85C protein, showing potential as lead compounds in TB drug development **[2].**

The discovery of these inhibitors has been facilitated by detailed structural analyses of the Ag85C protein **[11].** Understanding the precise structural dynamics of Ag85C, particularly how it interacts with various inhibitors, has been key to developing drugs that can effectively target this protein **[5;26].** Recent studies have explored the possible binding mechanisms of various compounds to Ag85C protein revealing some of the critical insights into how these interactions can be optimized for greater efficacy. For example, structural and dynamic studies have shown that certain mutations in Ag85C can significantly alter its binding properties, affect the protein’s function and its potential as a drug target **[15**].

In this study, we aim to further explore the potential of Ag85C as a target for TB drug discovery. Using *in-silico* approach, we will investigate the structural studies of Ag85C in complex with newly identified inhibitors. Our goal is to assess their binding affinities, stability, and potential as lead compounds in the fight against MDR and XDR-TB. Therefore by integrating computational simulations, we hope to contribute to the development of new, effective treatments that can overcome the challenges posed by drug-resistant TB.

## Materials and Methods

### Preparation of the Ligands

In this study the known ligand series which include Delamanid, Pretomanid, Benzothiazinones, Bedaquiline, Cyclipostins, and SQ109 were acquired from the respective literatures listed in the **Table 1** and illustrated in **Fig 1**. These ligands are prepared using Auto Dock 4.2 and LigPrep from Maestro for comparative studies **[31; 38; 16; 44**]. The structures were acquired from the PubChem database and converted into PDBQT format using Auto Dock Tools **[13].** This was achieved by integrating Gasteiger charges, merging non-polar hydrogens, and specifying rotatable bonds. LigPrep, a feature of the Maestro software, was utilized to generate 3D structures with optimized energy levels. The software utilized the OPLS4 force field to enhance the shape and structure of the molecules. Additionally, it assured that the molecules were appropriately ionized at a pH level consistent with physiological conditions. Later, these ligands were used to perform molecular docking studies to predict their interactions with the target protein Ag85C.

**Fig 1:**
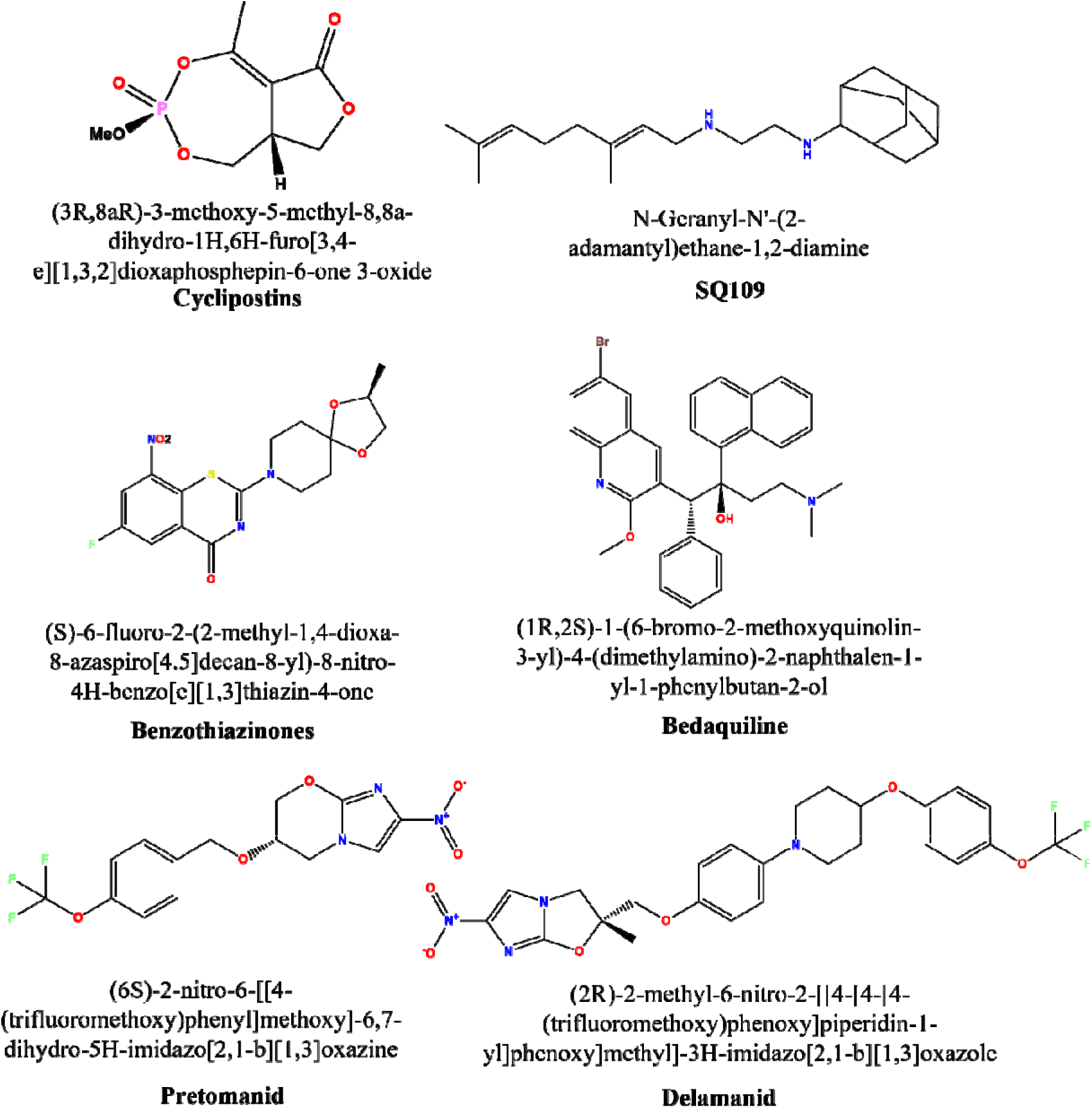
Systematic study of the target molecules for the development of active pharmacophore for the Mbt Ag85C from the selected target molecules.

**Table 1:**
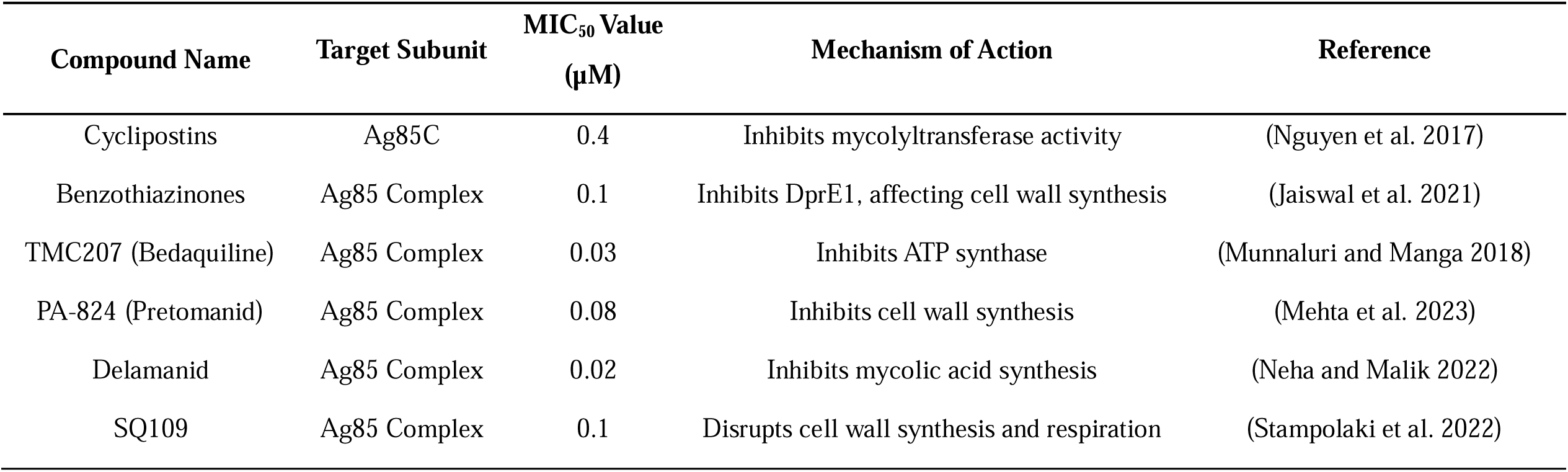
Target molecules available based on the available literature.

### Preparation of the Protein

The protein conformation of Ag85C (PDB ID: 5OCJ) was obtained from the RCSB Protein Data Bank **[26; 41**]. The function of this structure in the transport of mycolic acid is essential, as it plays a critical role in the formation of the cell wall of *Mtb* (refer the supporting information **Fig S1**). The protein preparation involves using Discovery Studio from BIOVIA to remove water molecules, add any missing hydrogen atoms, and enhance the geometry of the structure. Furthermore, the protein was pre-processed using the Protein Preparation Wizard in Maestro. This process entailed the allocation of bond ordering, insertion of hydrogen atoms, optimization of hydrogen bonds, and the execution of constrained minimization utilizing the OPLS4 force field. The protein structures were utilized for docking studies with the produced ligands.

### Molecular Docking

The protein prepared and prepared ligands (Delamanid, Pretomanid, Benzothiazinones, Bedaquiline, Cyclipostins, and SQ109) were used for molecular docking. The protein and ligands were converted to PDBQT (“.pdbqt”) format using AutoDock 4.0, and docking simulations were conducted to predict the binding affinities (in ΔG in Kcal/Mol) and interactions (dissociation constants (ki) in Molarity) (Vyshnava et al. 2020b). The grid box was configured to encompass the active site of Ag85C (the functional coordinates are x:-5.560321; y:-22.763250; z:-32.883321 with Area in: 11.600 Å), and the Lamarckian Genetic Algorithm was employed for docking. The ligands were docked into the generated protein structure using Maestro’s Ligand Docking module. The Glide docking tool utilized the extra precision (XP) mode to forecast binding modes and interactions. A comparative analysis was conducted to assess the relative interactions and binding affinities of the ligands with Ag85C by comparing the findings obtained from both AutoDock 4.0 and Maestro.

### Pharmacophore development

The protein and the ligands were subjected to molecular docking using AutoDock 4.0 and Maestro’s Ligand Docking module. The compound with the highest binding affinity and interaction profile was determined as the top hit. Subsequently, the leading hit was utilized to construct a pharmacophore model with Phase in Maestro (Schaller et al. 2020; Vrontaki and Kolocouris 2018). An analysis of the docking results was conducted to uncover crucial hydrogen bonds (HB), hydrophobic interactions (HI), and other important contacts between the ligand and the active site of Ag85C. The Phase module in Maestro was used to extract critical interaction properties, including hydrogen bond donors (HD) and acceptors (HA), hydrophobic areas (H), and aromatic rings (R), from the top hit compound. The aforementioned characteristics were employed in the preparation of a pharmacophore model, which accurately depicts the crucial interactions necessary for achieving both strong binding affinity and specificity towards Ag85C. The pharmacophore model was validated by a comparison with known active and inactive compounds. The model was refined to verify its accuracy in predicting the binding potential of related molecules.

### Screening of library

In order to find possible antituberculosis substances, we employed the Antituberculosis Screening Libraries provided by Life Chemicals (LifeChemials). There were two main collections involved in the preparation. The Antituberculosis Focused Library was generated by conducting a 2D fingerprint similarity search against a reference collection of biologically active chemicals. This search yielded 4,200 distinct molecules that specifically target critical proteins of the TB-causing bacterium, *Mtb*. The second library, known as the Antituberculosis Docking Library, consists of 7,600 compounds that were found by virtual molecular docking against specific therapeutic targets for tuberculosis. The libraries underwent filtration based on drug-likeness and ADMET characteristics to ensure the identification of promising candidates for subsequent validation.

### Confirmation of structures

The compounds obtained from screening the selected targets were subsequently submitted to molecular docking to confirm and compare them with the most effective known compound identified in prior molecular docking studies. To confirm the structure, AutoDock 4.0 and Maestro were used to dock the molecules into the generated protein (PDB ID: 5OCJ). The docking results were observed and examined using Discovery Studio, MGL Tools, and PyMOL software. The interactions and affinities of the main bindings were evaluated to verify the potential efficacy of the new compounds against Ag85C. The comparison analysis carried out in this study ensured the dependability of the docking data and permitted the identification of highly promising medicine candidates.

### ADME analysis of the studied drugs

We employed an *in-silico* methodology to generate pharmacological targets and predict their ADME features by employing the SwissADME program **[8].** The selected chemicals’ molecular structures were submitted into SwissADME in SMILES, which generated predict for crucial pharmacokinetic characteristics such as absorption, distribution, metabolism, excretion, and toxicity (ADMET). Our focus was specifically on the calculation of lipophilicity (logP), topological polar surface area (tPSA), molecular weight, and the number of hydrogen bond providers and acceptors. We assessed the substrate selectivity of P-glycoprotein in order to anticipate potential interactions with efflux transporters. The thorough computational study allowed us to evaluate the suitability of the compounds for pharmacological use and their potential to be absorbed by the body, offering vital knowledge for future medication development.

## Results and Discussions

### Primary Docking of Selected Drugs on Ag85C

Delamanid and Pretomanid were identified as the top hits in the primary docking of selected drugs using Ag85C from *Mtb*. The interaction energies (ΔG) were –7.226 Kcal/mol and –8.58 Kcal/mol with dissociation constants (Ki) of 115.36 nM and 516.53 nM, respectively, based on AutoDock4.0. While the Glide docking scores were –7.226 and –6.984 for Delamanid and Pretomanid, respectively. Delamanid interacted with amino acids Ag85C B:ASP38, B:LEU40, B:ARG41, B:GLN43, B:ASN52, A:ALA165, B:GLY39, B:GLY48, A:ASP168, and B:TRP262 for the Delamanid and while Pretomanid interacted with A:ASP168, and B:PHE226, the details are listed in the **Table 2** and illustrated in **Fig 2**. The strong binding affinities of Delamanid and Pretomanid suggest their potential as effective inhibitors of Ag85C, which are consistent with the earlier reports [**21].** These interactions are significant as they highlight the crucial amino acids involved in the active site, which play a pivotal role in the inhibition of the enzyme. The effectiveness of Delamanid in treating multidrug-resistant tuberculosis (MDR-TB) is well-documented due to its inhibition of mycolic acid biosynthesis [**30].** Pretomanid bactericidal activity against both replicating and non-replicating *Mtb* further supports its inclusion as a top hit [**10].**

**Fig 2:**
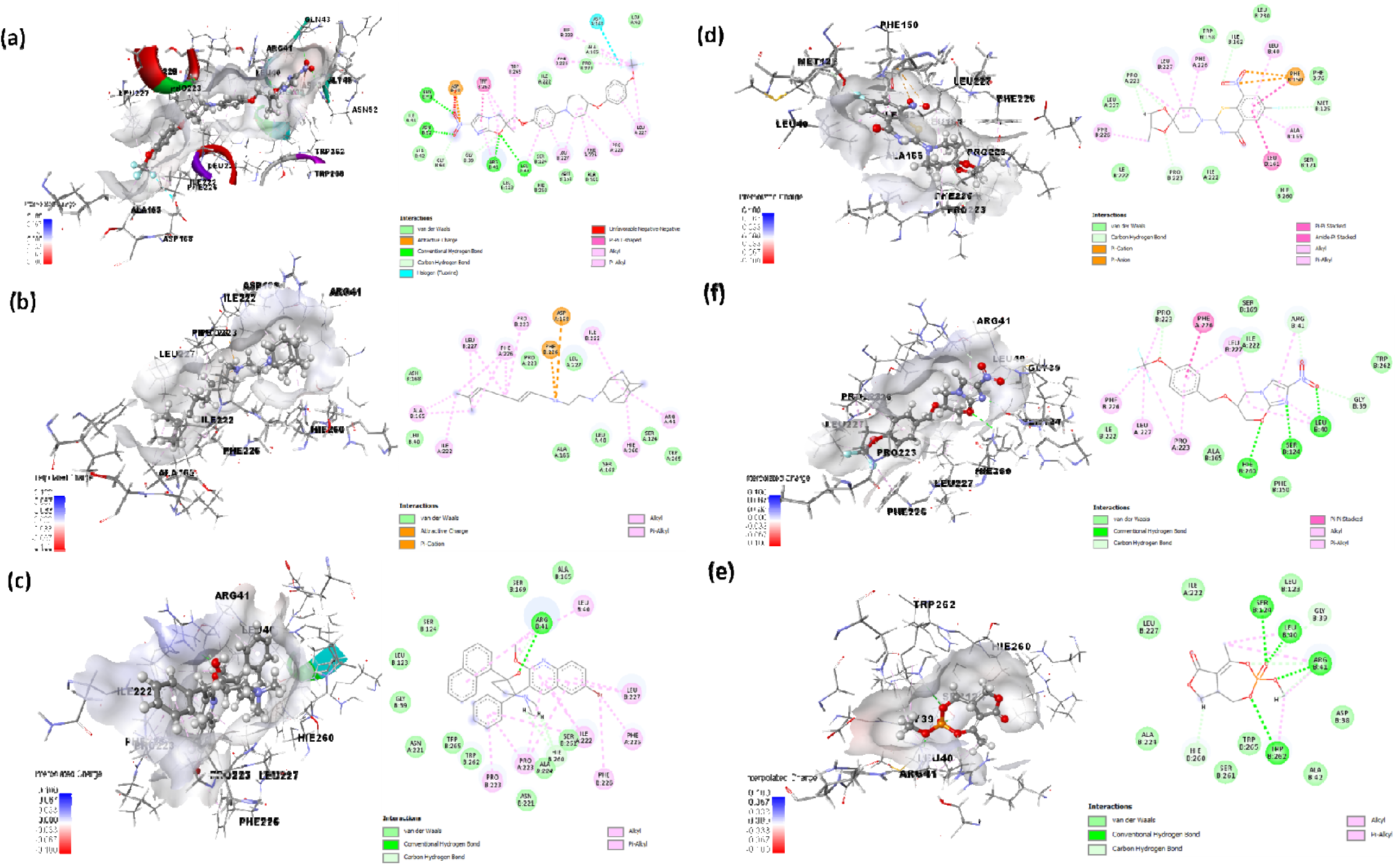
Molecular docking of the selected targeted compounds with the Mbt Ag85 complex. (a) Delamanid (b) SQ109 (c) Bedaquiline (d) Benzothiazinones (e) Pretomanid (f) Cyclipostins

**Table 2:** Molecular docking glide and Autodock 4.2 ligand receptor interactions for the targeted and screened drugs based on the top hits.

| Compounds of interest | Glide |  |  | AutoDock 4.0 |  | Interactive amino acids in the binding pockets |  |  |  |  |
| --- | --- | --- | --- | --- | --- | --- | --- | --- | --- | --- |
| | Docking score | XP GScore | Glide energy | Interaction energy $\Delta G$ (kcal/mol) | Dissociation constant $K_i$ (Molar) | Attractive charges | Hydrogen bonds (Conventional) | Hydrogen bonds (Carbon) | Halogen | Pi-Pi |
| Delamanid | -7.226 | -7.240 | -57.105 | -9.47 | 115.36 nM | B:ASP38 | B: LEU40<br>B: ARG41<br>B: GLN43<br>B: ASN52 | A:ALA165<br>B:GLY39<br>B:GLY48 | A:ASP168 | B:TRP262 |
| Pretomanid | -6.984 | -6.984 | -37.482 | -8.58 | 516.53 nM | A:ASP168<br>B:PHE226 | - | - | - | - |
| Benzothiazinones | -5.411 | -5.795 | -37.831 | -8.23 | 927.37 nM | - | B:ARG41 | B:ARG41<br>B:HIE260 | - | - |
| Bedaquiline | -4.465 | -4.470 | -33.997 | -8.10 | 1.15 μM | B:PHE150 | - | A:PRO223<br>B:MET125<br>B:ILE162<br>B:PRO223<br>A:PRO223 | - | B:PHE150<br>B:LEU161 |
| Cyclipostins | -3.322 | -3.322 | -26.641 | -6.68 | 12.75 μM | - | B:LEU40<br>B:SER124<br>B:HIE260 | B:GLY39<br>B:ARG41<br>B:PRO223 | - | A:PHE226 |
| SQ109 | -1.997 | -3.549 | -29.121 | -6.35 | 22.05 μM | - | B:LEU40<br>B:ARG41<br>B:SER124<br>B:TRP262 | B:GLY39<br>B:ARG41<br>B:SER124<br>B:HIE260<br>B:ARG41 | - | - |

These interactions are crucial for rational drug design as they provide insight into the structural requirements for effective binding, guiding the development of new inhibitors targeting Ag85C. The binding of Delamanid to multiple amino acids, including those involved in the stabilization of the enzyme’s active conformation, suggests that it may effectively block the catalytic activity of Ag85C. Pretomanid interaction with key residues such as A:ASP168, and B:PHE226 highlights the importance of these residues in the active site and their role in enzyme inhibition. The structural insights gained from these docking studies can inform the design of derivatives with enhanced binding affinities and specificities. Moreover, these findings align with previous research that emphasizes the importance of HB and HI in stabilizing ligand-receptor complexes. Understanding these interactions helps in designing drugs that not only bind effectively but also demonstrate high specificity, reducing potential off-target effects [**27**].

### Phase Pharmacophore Development

From the active and inactive forms of selected ligands, Bedaquiline and Delamanid were sorted as active, while Benzothiazinones, Cyclipostins, and Pretomanid were classified as inactive based on pIC50-µM values which are listed in **Table 3**. The phase pharmacophore method identified key pharmacophore features for active alignment, including A2, A3 as acceptors, R11 as a ring structure, and H6 and H9 as hydrophobic groups, which are shown in **Fig 3**. The development of a robust pharmacophore model is critical for virtual screening and drug design. The pharmacophore features identified in this study are consistent with known structural requirements for effective enzyme inhibition. The high sensitivity and selectivity of the model in distinguishing between active and inactive compounds validate its use in screening large libraries for potential drug candidates.

**Fig 3:**
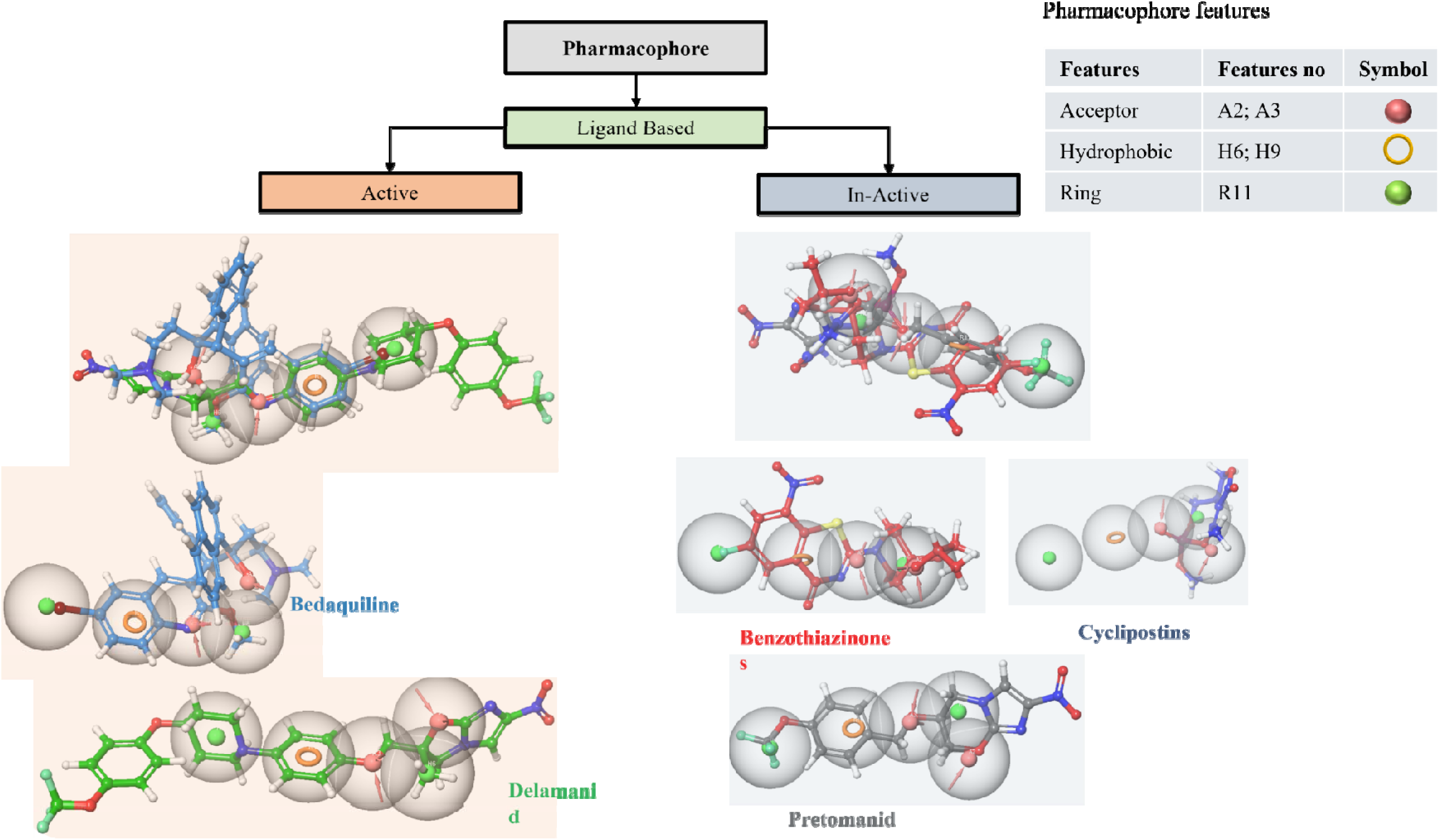
Pharmacophore development from the ligands based on the functional activity.

**Table 3:** Pharmacophore model Development by pIC_50_ features.

| Known Compounds | IC<sub>50</sub> (μM) | pIC<sub>50</sub> (μM) | Selected Features |
| --- | --- | --- | --- |
| Bedaquiline | 0.03 | 7.522879 | Active |
| Benzothiazinones | 0.1 | 7.000000 | In-Active |
| Cyclipostins | 0.4 | 6.397940 | In-Active |
| Delamanid | 0.02 | 7.698970 | Active |
| Pretomanid | 0.08 | 7.096910 | In-Active |
| SQ109 | 0.1 | 7.000000 | In-Active |

**Note:** pIC<sub>50</sub> μM= -1og Exp (PIC<sub>50</sub> μM)

**Table 4:**
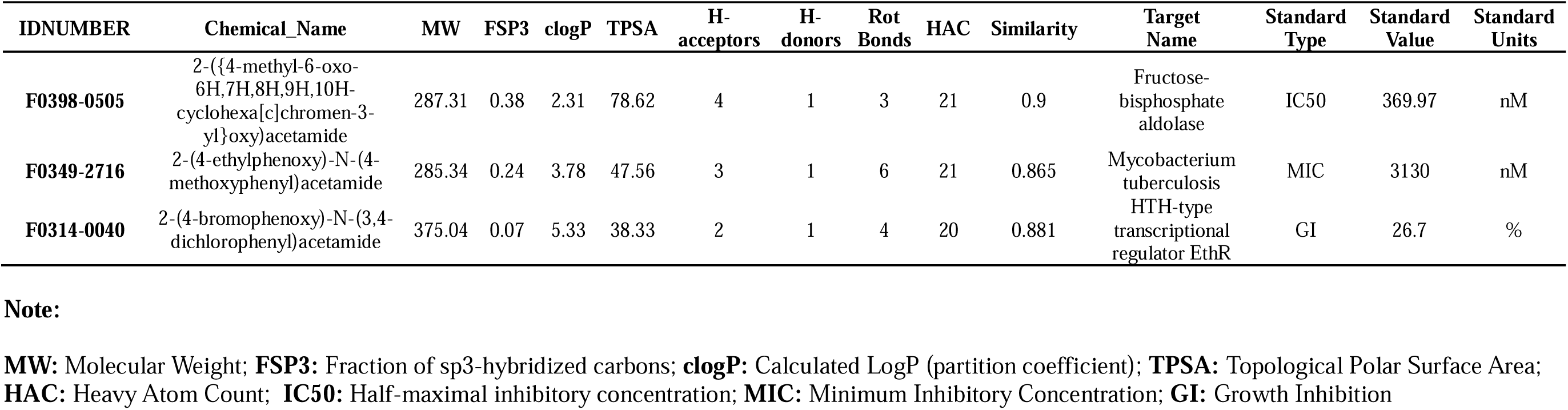
Top targets for the developed pharmacophore.

**Table 5:**
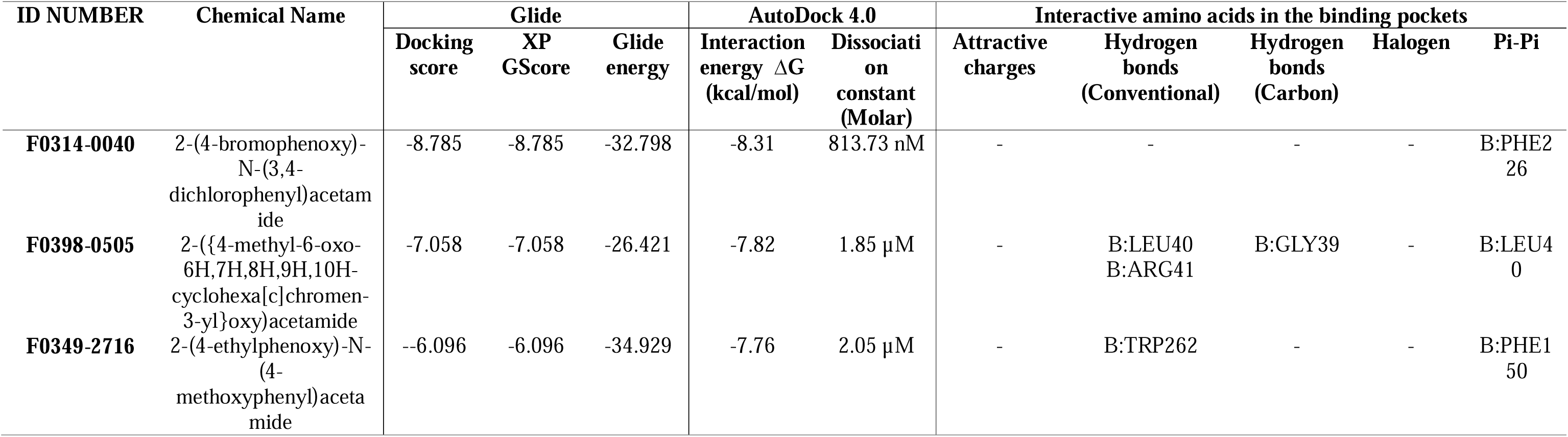
Molecular docking glide and AutoDock 4.2 ligand receptor interactions for the screened top hit compounds.

Previous studies have emphasized the importance of pharmacophore modeling in identifying key molecular interactions necessary for binding and activity. The identified features, such as HB acceptors and HB donors, R structures, and H groups, are essential for binding to the active site of Ag85C. This rational approach allows for the identification of new compounds that can effectively target the Ag85C enzyme. Pharmacophore modelling provides a virtual template for the optimal spatial arrangement of functional groups necessary for molecular recognition at the binding site. The ability to predict and validate pharmacophore models ensures that only the most promising candidates are selected for further development [**34; 40**].

### Identification of New Drug Candidates

Screening of 4254 antituberculosis compounds acquired from Life Chemicals and the ChEMBL open-source database identified three top hits fitting the developed pharmacophore: F0314-0040 (CHEMBL2164321), F0398-0505 (CHEMBL1608557), and F0349-2716 (CHEMBL3637857), which are depicted in **Fig 4** and **Fig 5**. The identification of these compounds demonstrates the utility of the pharmacophore model in discovering novel drug candidates. The use of high-throughput screening and large compound libraries enhances the chances of finding potent inhibitors. The selected compounds were found to possess structural features compatible with the pharmacophore model, indicating their potential effectiveness in inhibiting Ag85C.

**Fig 4:**
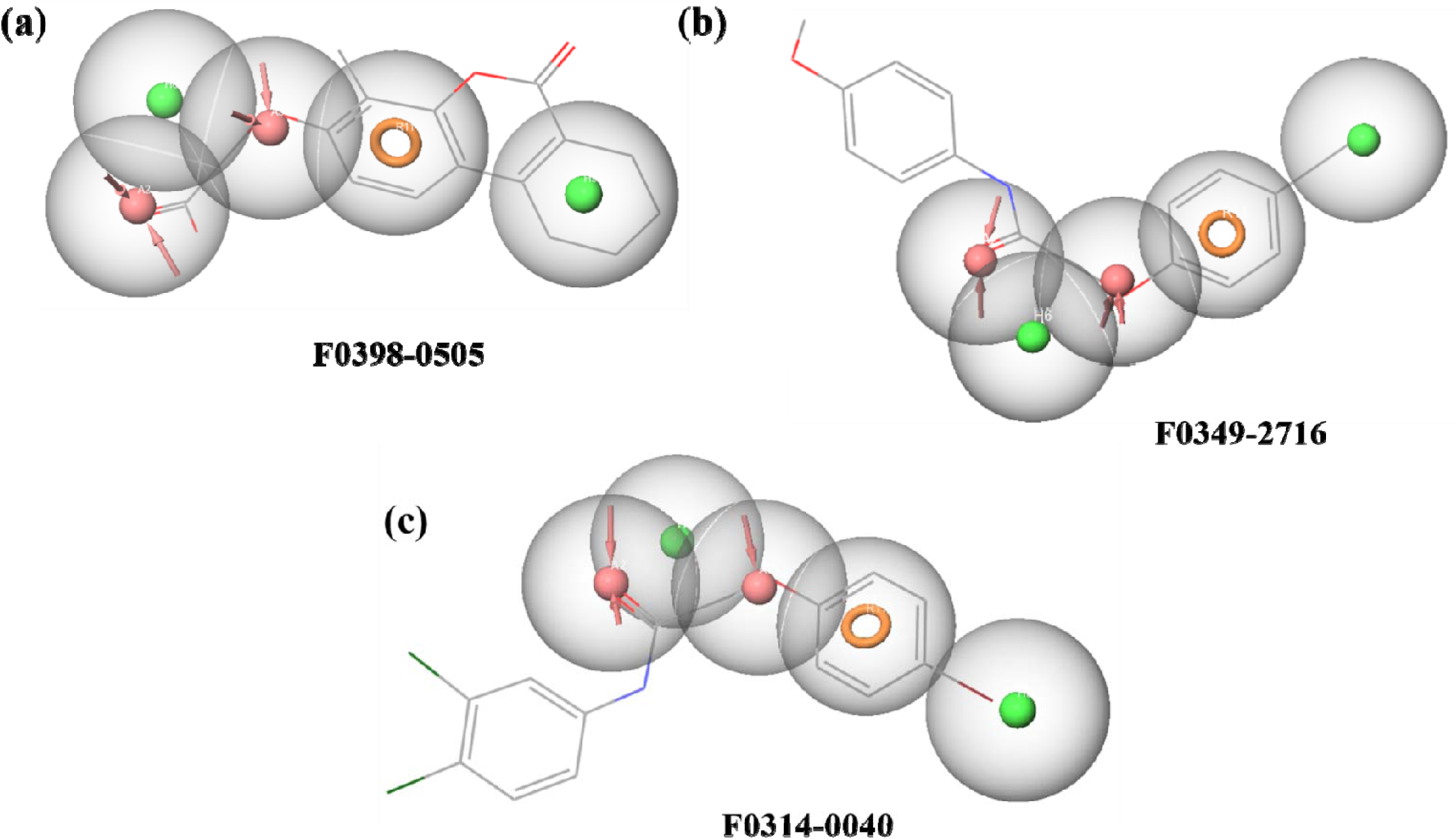
Pharmacophore development from the ligands screened drugs.

**Fig 5:**
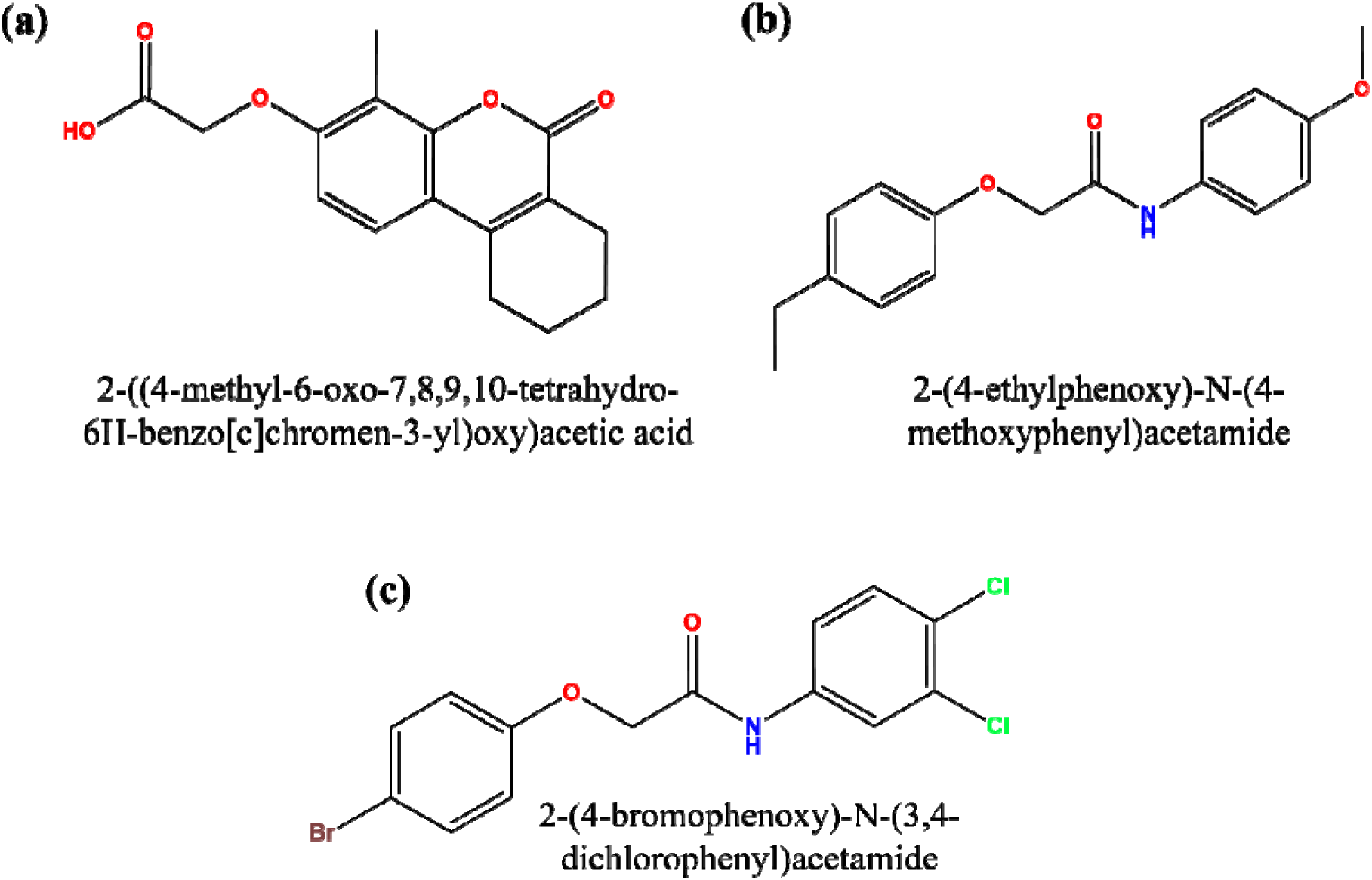
Top hits of the screened based on the Pharmacophore.

This approach aligns with the strategies used in previous drug discovery efforts where large libraries are screened to identify hits that fit a validated pharmacophore model **[24; 20**]. The ability to identify new compounds with high binding affinities and appropriate pharmacophore features is crucial for the development of new antituberculosis drugs. These findings highlight the importance of integrating pharmacophore modeling with virtual screening to streamline the drug discovery process. The success of this screening also underscores the importance of accessing diverse chemical libraries. By using databases such as ChEMBL Life Chemicals (drug library build based on the ChEMBL registered compounds), researchers can tap into a wide array of chemical entities, increasing the likelihood of identifying effective drug candidates. This strategy ensures that potential inhibitors are not overlooked due to limited chemical diversity **[39; 29**].

### Major Docking of New Drug Candidates

F0314-0040 (CHEMBL2164321) and F0398-0505 (CHEMBL1608557) were the top hits in major docking against the Ag85C active site. Interaction energies (ΔG) were –8.31 Kcal/mol and –7.82 Kcal/mol with dissociation constants (Ki) of 813.73 nM and 1.85 µM, respectively. Glide docking scores were –8.785 and –7.058, respectively. Key interactions included amino acid B:PHE226 which are depicted in **Fig 6**. The major docking results underscore the importance of specific amino acid interactions in achieving strong binding affinities. The identified compounds exhibited competitive binding energies and dissociation constants, indicating their potential as effective inhibitors. The interactions with key amino acids B:GLY39, B:LEU40, B:ARG41 are crucial as they contribute to the stability and efficacy of the binding.

**Fig 6:**
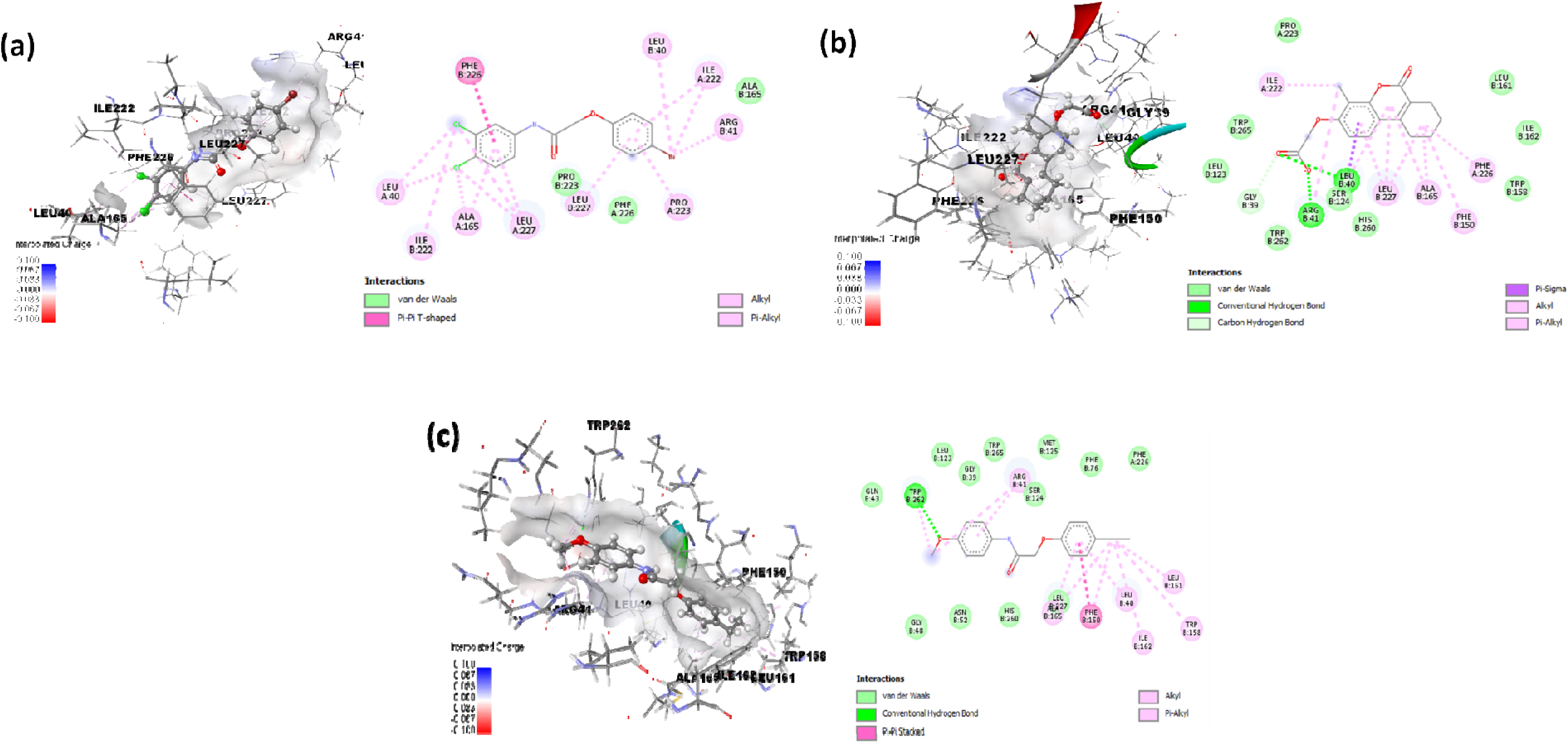
Molecular docking of the Pharmacophore based screened ligands. (a) F0398-0505 (b) F0349-2716 (c) F0314-0040

The detailed understanding of these interactions provides a basis for rational drug design, allowing for the optimization of compounds to enhance their binding affinities and inhibitory activities. The strong binding affinities observed for F0314-0040 and F0398-0505 suggest that these compounds are promising candidates for further development and testing.

The docking studies also provide insights into the structural aspects of drug design, emphasizing the need to consider both the binding energy and the specific interactions with the active site residues. The binding modes observed in this study can inform the design of analogs with improved properties, such as increased potency and selectivity. The use of computational methods to predict binding interactions and optimize drug candidates is a powerful approach in the early stages of drug discovery.

Moreover, the interaction of these compounds with the key residues involved in the enzyme’s catalytic activity supports their potential as inhibitors. The ability to disrupt the function of Ag85C by binding to these critical sites underpins the rationale for targeting these residues in drug design. This strategy is aligned with the principles of structure-based drug design, where detailed knowledge of the target structure guides the development of more effective therapeutic agents [**32**].

### ADMET analysis

The ADMET (Absorption, Distribution, Metabolism, Excretion and Toxicity) properties of the molecules of interest were evaluated using SwissADME. Delamanid, SQ109, Bedaquiline, Benzothiazinones, Pretomanid, Cyclipostins, F0314-0040, F0398-0505, and F0349-2716 were assessed for Druglikeness and pharmacokinetics. Delamanid and Pretomanid exhibited favourable Druglikeness with high consensus LogP values and acceptable bioavailability scores, suggesting their potential as promising candidates for further development, these properties are listed in the **Table S1** and **Table S2** (refer the supporting information). Their low number of rule violations indicates a favourable balance between hydrophilicity and lipophilicity, which is essential for drug absorption and distribution. Conversely, Bedaquiline, despite its high binding affinity, had multiple rule violations and a lower bioavailability score, which could limit its effectiveness due to poor absorption and potential off-target effects. The high Log P value of Bedaquiline indicates strong lipophilicity, potentially leading to extensive distribution into fatty tissues and associated toxicity. SQ109 and Benzothiazinones showed favourable Druglikeness profiles with minimal rule violations and good bioavailability scores, supporting their potential as viable drug candidates. The novel compounds F0314-0040, F0398-0505, and F0349-2716 demonstrated promising Druglikeness with no rule violations and high bioavailability scores, highlighting their potential for further investigation, which are listed in **Table S3** (refer the supporting information).

The pharmacokinetics analysis revealed that Delamanid exhibited low gastrointestinal absorption but significant cytochrome P450 enzyme inhibition, indicating potential challenges in oral administration and possible drug-drug interactions. These properties need to be optimized for a better pharmacokinetic profile. SQ109 and the novel compounds F0314-0040, F0398-0505, and F0349-2716 demonstrated high gastrointestinal absorption and BBB permeability, indicating potential effectiveness in systemic and central nervous system infections. However, the inhibition of multiple cytochrome P450 enzymes by F0314-0040 and F0349-2716 suggests caution regarding metabolic interactions, which are shown in the **Fig S2** (refer the supporting information). Bedaquiline’s low GI absorption and high interaction with P-glycoprotein and cytochrome P450 enzymes highlight the need for formulation strategies to enhance its bioavailability and reduce potential side effects. Benzothiazinones and Pretomanid exhibited high GI absorption and selective enzyme inhibition, making them suitable for oral administration with manageable metabolic profiles. Cyclipostins’ high GI absorption and lack of significant enzyme inhibition suggest a favourable pharmacokinetic profile, although its lower Log P value indicates a need to balance hydrophilicity for effective distribution.

## Conclusions

The integration of docking, pharmacophore modelling, and ADME/pharmacokinetics analysis identified Delamanid and Pretomanid as effective inhibitors of the Ag85C enzyme, with strong binding affinities and key interactions with the active site. The pharmacophore model successfully identified three new drug candidates with promising antituberculosis activity. These findings support further *in vitro* and *in vivo* testing to confirm the efficacy and safety of the identified compounds, enhancing the drug discovery process for tuberculosis treatment.

## Author contributions

Dr. Anuradha and Dr. Chitta Suresh Kumar accepted & approved for carrying out the work and verified the data by analysis. Mr. Katewad Raghavendra Singh carried out the in-silico work and manuscript preparation.

## Declarations

### Ethics approval and consent to participate

Not Applicable as no animals were used to carried out this study.

## Consent for publication

Not Applicable

## Data availability

The authors declare that all other data supporting in this study are available within the paper and its supplementary information.

