## Supplementary info for "*In-silico* Exploration of Drug Leads Targeting Mycobacterial Ag85C"

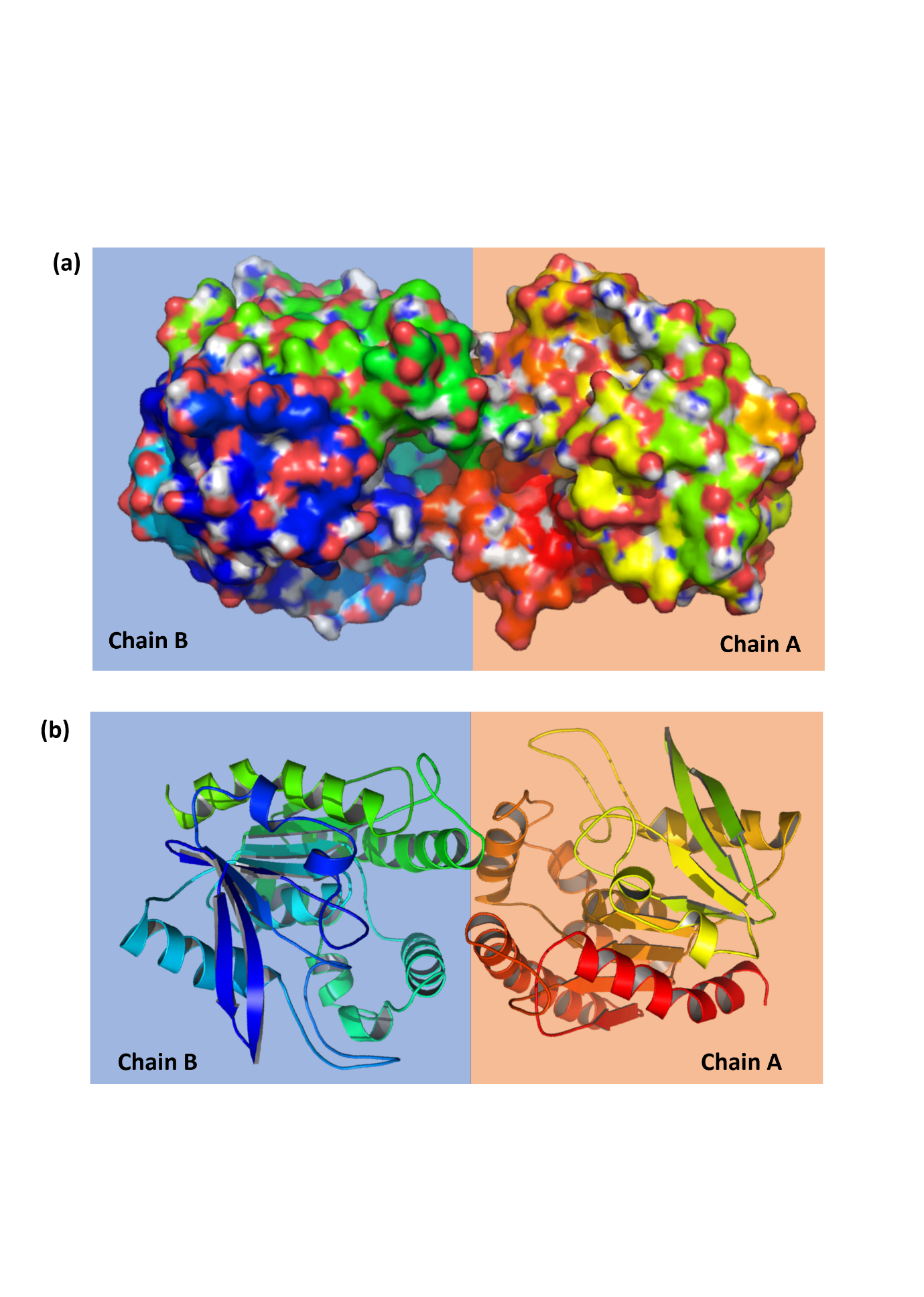

**Fig S1:** Molecular structure of Ag85C (PDB ID:5OCJ) (a) Surface model with active site pockets (b) Ribbon model showing the helix and the sheets

**Table S1:** Druglikeliness of molecules of interest List -I

| **Molecule** | **iLOGP** | **XLOGP3** | **WLOGP** | **MLOGP** | **Silicos-IT**  **Log P** | **Consensus**  **Log P** |
| --- | --- | --- | --- | --- | --- | --- |
| Delamanid | 3.71 | 5.63 | 5.85 | 2.59 | 1.55 | 3.87 |
| SQ109 | 4.91 | 5.11 | 4.68 | 4.26 | 4.61 | 4.71 |
| Bedaquiline | 4.74 | 7.21 | 7.02 | 5.08 | 7.18 | 6.25 |
| Benzothiazinones | 2.57 | 2.13 | 2.47 | 1.82 | 1.65 | 2.13 |
| Pretomanid | 2.52 | 2.78 | 3.78 | 1.12 | 0.09 | 2.06 |
| Cyclipostins | 1.73 | -0.25 | 1.23 | -0.39 | 0.93 | 0.65 |
| F0314-0040 | 2.77 | 4.99 | 4.58 | 4.07 | 4.43 | 4.17 |
| F0398-0505 | 2.2 | 2.78 | 2.44 | 2.01 | 3.59 | 2.61 |
| F0349-2716 | 2.83 | 3.5 | 3.08 | 2.56 | 3.41 | 3.08 |

**Table S2:** Druglikeliness of molecules of interest List -II

| **Molecule** | **Lipinski**  **#violations** | **Ghose**  **#violations** | **Veber**  **#violations** | **Egan**  **#violations** | **Muegge**  **#violations** | **Bioavailability**  **Score** |
| --- | --- | --- | --- | --- | --- | --- |
| Delamanid | 1 | 3 | 0 | 0 | 1 | 0.55 |
| SQ109 | 1 | 0 | 0 | 0 | 1 | 0.55 |
| Bedaquiline | 2 | 3 | 0 | 1 | 1 | 0.17 |
| Benzothiazinones | 0 | 0 | 0 | 0 | 0 | 0.55 |
| Pretomanid | 0 | 0 | 0 | 0 | 0 | 0.55 |
| Cyclipostins | 0 | 0 | 0 | 0 | 0 | 0.56 |
| F0314-0040 | 0 | 0 | 0 | 0 | 0 | 0.55 |
| F0398-0505 | 0 | 0 | 0 | 0 | 0 | 0.56 |
| F0349-2716 | 0 | 0 | 0 | 0 | 0 | 0.55 |

**Note:**

Lipinski #violations: MW ≤ 500; Lipophilicity property octanol-water partition coefficient (MLOGP) ≤ 4.15; N or O ≤ 10; NH or OH ≤ 5. Ghose #violations: 160 ≤ MW ≤ 480; -0.4 ≤ WLOP ≤ 5.6; 40 ≤ Molecular refractivity (MR) ≤ 130; 20 ≤ atoms ≤ 70. Veber #violations: Rotatable bonds ≤ 10; Topological polar surface area (TPSA) ≤ 140. Egan #violations: WLOGP ≤ 5.88; TPSA ≤ 131.6. Muegge #violations: 200 ≤ MW ≤ 600; -2 ≤ XLOP ≤ 5; TPSA ≤ 150; Num. rings ≤ 7; Num.Carbons ≤ 4; Num RB ≤ 15; H-Bonds acc ≤ 10; H-Bonds don ≤ 5. Bioavailability Score: Probability factor F > 10% in rats

**Table S3:** Pharmacokinetics of molecules of interest

| **Molecule** | **GI**  **absorption** | **BBB**  **permeant** | **Pgp**  **substrate** | **CYP1A2**  **inhibitor** | **CYP2C19**  **inhibitor** | **CYP2C9**  **inhibitor** | **CYP2D6**  **inhibitor** | **CYP3A4**  **inhibitor** | **log Kp**  **(cm/s)** |
| --- | --- | --- | --- | --- | --- | --- | --- | --- | --- |
| Delamanid | Low | No | No | No | Yes | Yes | Yes | Yes | -5.56 |
| SQ109 | High | Yes | No | No | No | No | Yes | No | -4.69 |
| Bedaquiline | Low | No | Yes | Yes | No | No | Yes | Yes | -4.57 |
| Benzothiazinones | High | No | Yes | Yes | Yes | Yes | No | No | -7.11 |
| Pretomanid | High | No | No | Yes | Yes | Yes | No | No | -6.52 |
| Cyclipostins | High | No | No | No | No | No | No | No | -7.91 |
| F0314-0040 | High | Yes | No | Yes | Yes | Yes | Yes | No | -5.04 |
| F0398-0505 | High | Yes | No | Yes | No | No | No | No | -6.08 |
| F0349-2716 | High | Yes | No | Yes | Yes | Yes | Yes | No | -5.56 |

**Note:**

GIA- Gastrointestinal absorption; BBBP-Blood brain barrier permeant; Pgps- p Glycoprotein substrate; CYP1A2- Cytochrome P450 A2 inhibitor; CYP2C19- Cytochrome P450 2C19 inhibitor; CYP2C9- Cytochrome P450 2C9 inhibitors; CYP2D6- Cytochrome P450 2D6 inhibitor; CYP3A4- Cytochrome P450 3A4; log Kp (cm/s)- Skin Permeation

| **Compound** | **Molecule No** |
| --- | --- |
| **Cyclipostins** | Molecule 1 |
| **SQ109** | Molecule 2 |
| **Benzothiazinones** | Molecule 3 |
| **Bedaquiline** | Molecule 4 |
| **Pretomanid** | Molecule 5 |
| **Delamanid** | Molecule 6 |
| **F0398-0505** | Molecule 7 |
| **F0349-2716** | Molecule 8 |
| **F0314-0040** | Molecule 9 |

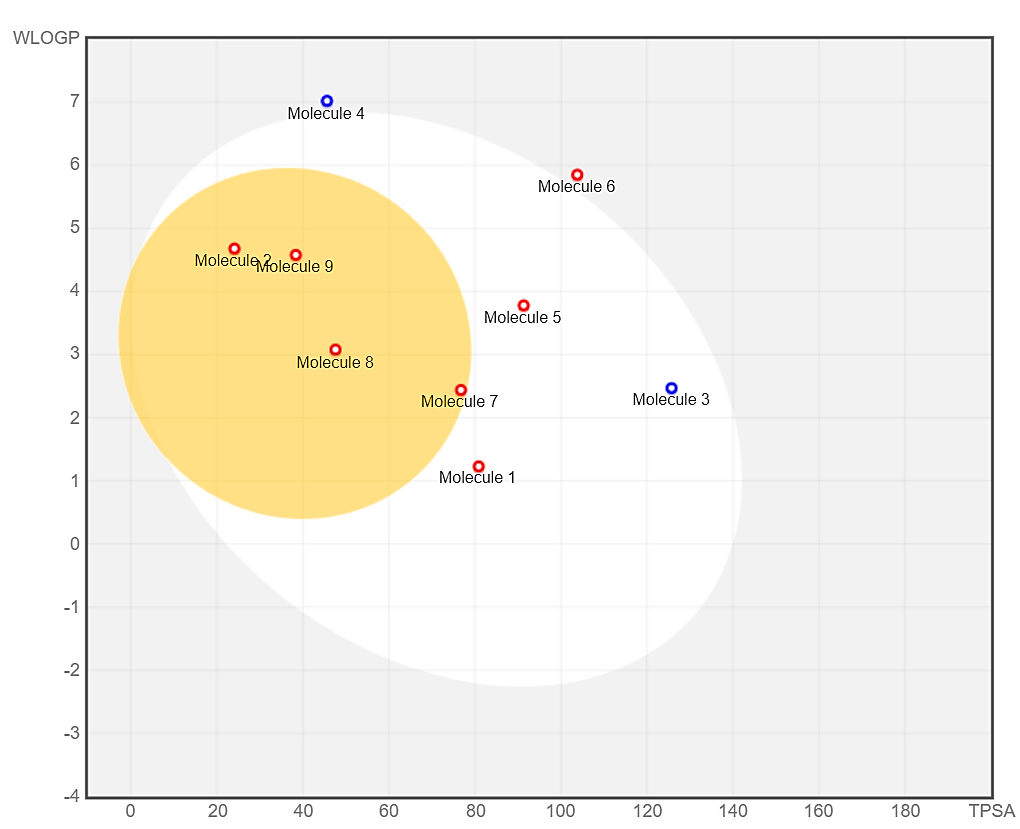

**Fig S2:** BOILED-Egg model for the known and screened targets of interest

**Note:** Compounds within the white ellipse have a high likelihood of being passively absorbed through the gastrointestinal tract. Those within the yellow ellipse, or "yolk," are predicted to permeate the blood-brain barrier and reach the central nervous system. These regions can overlap, indicating compounds with both properties. Compounds outside these ellipses fall into the grey zone, suggesting they are less likely to be well absorbed or penetrate the BBB.

Compounds indicated in blue are predicted substrates of P-glycoprotein (PGP+), meaning they are likely to be actively expelled from the brain or back into the gastrointestinal lumen. Conversely, compounds shown in red are predicted non-substrates of P-glycoprotein (PGP-).
